# Targeted development of diagnostic SNP markers for resistance to Fusarium wilt race 4 in Upland cotton (*Gossypium hirsutum*)

**DOI:** 10.1101/2023.03.06.531315

**Authors:** Jinfa Zhang, Yi Zhu, Terry Wheeler, Jane K Dever, Kater Hake, Kaitlyn Bissonnette

**Affiliations:** Department of Plant and Environmental Sciences, New Mexico State University, Las Cruces, NM 88003, USA; Texas A&M AgriLife Research, 1102 E. Drew Street, Lubbock, TX 79403, USA; Cotton Incorporated, 6399 Weston Parkway, Cary, NC 27513, USA

**Keywords:** Upland cotton, Fusarium wilt, Race 4, Resistance, SNPs

## Abstract

Fusarium wilt caused by the soil-borne fungus *Fusarium oxysporum* f. sp. *vasinfectum* (FOV) race 4 (FOV4) has become one of the most important emerging diseases in US cotton production. Numerous QTLs have been reported for resistance to FOV; however, no major FOV4-resistance QTL or gene has been identified and used in breeding Upland cotton (*Gossypium hirsutum*) for FOV4 resistance. In this study, a panel of 223 Chinese Upland cotton accessions was evaluated for FOV4 resistance based on seedling mortality rate (MR) and stem and root vascular discoloration (SVD and RVD). SNP markers were developed based on targeted genome sequencing using AgriPlex Genomics. The chromosome region at 2.130-2.292 Mb on D03 was significantly correlated with both SVD and RVD but not with MR. Based on the two most significant SNP markers, accessions homozygous for AA or TT SNP genotype averaged significantly lower SVD (0.88 vs. 2.54) and RVD (1.46 vs. 3.02) than those homozygous for CC or GG SNP genotype. The results suggested that a gene or genes within the region conferred resistance to vascular discoloration caused by FOV4. The Chinese Upland accessions had 37.22% homozygous AA or TT SNP genotype and 11.66% heterozygous AC or TG SNP genotype, while 32 US elite public breeding lines all had the CC or GG SNP genotype. Among 463 obsolete US Upland accessions, only 0.86% possessed the AA or TT SNP genotype. This study, for the first time, has developed diagnostic SNPs for marker-assisted selection and identified FOV4-resistant Upland germplasms with the SNPs.

## Introduction

Fusarium wilt, caused by the soil- and seed-borne fungal pathogen *Fusarium oxysporum* f. sp. *vasinfectum* Atk. Sny & Hans (FOV), is one of the most destructive diseases in cotton (*Gossypium* spp.) worldwide. It causes significant yield losses ranging 0.19 to 1.36% across the US Cotton Belt (Blasingame and Patel 2013) and as high as 2.5% in California (Lawrence et al. 2021). The FOV pathogen penetrates cotton roots through epidermal cells and then invades and colonizes the vascular system. The infection produces toxic compounds and prevents absorption and transport of water and nutrients, causing vascular discoloration, plant wilting, stunted growth, leaf chlorosis and necrosis, defoliation, and plant death (Zhang et al. 2015; Sanogo and Zhang 2016). In addition to a virulent Australian strain, eight FOV pathogenic races (designated races 1 to 8) have been reported in the world, based on pathogenicity tests and limited DNA marker or sequence analysis (Cianchetta et al. 2015; Davis et al. 2006). They are divided into four groups: Group 1-races 1, 2, and 6; Group 2-races 3 and 5 (both are currently indistinguishable based on DNA markers); Group 3-races 4 and 7 [(both belong to the same FOV vegetative compatibility group (Bell et al. 2019) and are also currently indistinguishable based on DNA markers and limited gene sequences (Zhu et al. 2021a)]; and Group 4-race 8. In the US, races 1, 2, 3, 4 and 8, and several new FOV genotypes are identified (Holmes et al. 2009; Davis et al. 2006; Cianchetta and Davis 2015; Cianchetta et al. 2015). Races 1 and 2 require interactions with nematodes including primarily root-knot nematodes [*Meloidogyne incognita* (Kofoid & White] Chitwood)] for a successful infection under field conditions, and they are most widely distributed across the US Cotton Belt. However, FOV race 4 (FOV4), originally reported in India (Armstrong and Armstrong, 1960), is the most destructive race in the US, because it causes an early season disease with seedling wilt and death immediately after emergence in the absence of nematodes and subsequently significant yield losses in the field (Zhang et al. 2020c).

In the US, FOV4 was first identified in California in 2001 (Kim et al. 2005) and has become a significant production problem in both Pima (*G. barbadense* L.) and Upland (*G. hirsutum* L.) cotton in California (Diaz et al. 2021). It has since been identified in far-west Texas (Halpern et al. 2018; Bell et al. 2019; Diaz et al. 2021; Zhu et al. 2019, 2021a; Wagner et al. 2022) and southern New Mexico (Zhang et al. 2020d; Zhu et al. 2020, 2021b). Its potential to expand to other cotton production regions is a cause for concern. Four different FOV4 genotypes were recently identified based on the presence of different insertions in the *PHO* gene encoding for phosphate permease (Bell et al. 2019); however, the relationship between the four FOV4 genotypes and virulence is currently unclear. The frequencies of the four FOV4 genotypes differed among Texas, California, New Mexico, China, and India (Bell et al. 2019; Zhu et al. 2021a). Because of its soil-borne nature, saprophytic growth, and long-term survival ability, controlling the disease through chemical, rotational, and physical methods is extremely difficult. The most cost-effective method to manage FOV is through breeding and utilizing resistant cotton cultivars, as exemplified by the extensive and successful breeding effort for FOV7 resistance in Upland cotton in China (Zhang et al. 2015; Sanogo and Zhang 2016; Zhang 2018) and for FOV4 resistance in herbicide-tolerant Pima cotton such as PHY 802 RF, PHY 805 RF, PHY 811 RF, and PHY 841 RF with FOV4 resistance derived from Pima S-6 in the US.

Resistance to FOV4 was determined by two complementary dominant genes and a third inhibitory gene in *G. arboreum* L. and one dominant gene in *G. herbaceum* L. in India (cited by Smith and Dick 1960). *G. arboreum* germplasm lines with resistance to FOV4 were recently identified in the US cotton germplasm collection (Abdelraheem et al. 2021), but no genetic studies in resistance have been reported. Based on a genome-wide association study (GWAS) using 215 Chinese *G. arboreum* accessions with 1.4 million resequencing-based single nucleotide polymorphic (SNP) markers, *Ga11G2353* encoding for a glutathione S-transferase (GaGSTF9) was identified to be the candidate gene for a major-effect quantitative trait locus (QTL) on chromosome A11 (within a 1.0 Mb interval in the 102.5-103.5 Mb region) for resistance to race 7, i.e., FOV7 (Du et al. 2018). However, the resistance gene(s) against FOV4 or FOV7 has not been transferred into Upland cotton, and no resistance QTL on A11 for FOV4 or FOV7 resistance has been identified in tetraploid cotton (Zhu et al. 2022).

In *G. barbadense*, a major resistance QTL (designated *FOV4*) conferring resistance to FOV4 was identified in resistant Pima S-6, which was initially mapped to D02 through a linkage analysis using simple sequence repeat (SSR) markers (Ulloa et al. 2013). However, the DNA sequences for the anchoring SSR markers BNL0834 and MUSS354 are in fact on D03 (c17) at the 0.78 and 2.25 Mb regions, respectively, based on the recently sequenced *G. hirsutum* (e.g., TM-1) and *G. barbadense* (e.g., H7124 and 3-79) genomes (Hu et al. 2019; Wang et al. 2019; www.cottongen.org). The D03 location (at 1.15-2.29 Mb) of the resistance gene for FOV4 in *G. barbadense* was later confirmed using an F2 population involving Pima PHY 800 (with resistance derived from Pima S-6) and SNP markers (https://patentimages.storage.googleapis.com/20/77/cd/e066a5b4503386/WO2020139756A1.pdf). However, no candidate gene(s) for the FOV4-resistance QTL has been reported. Interestingly, a major resistance QTL to FOV7 on D03 at 1.4-1.6 Mb with two candidate genes (*Gbar_D03G001430* and *Gbar_D03G001910*) was recently reported in *G. barbadense* based on GWAS of 336 accessions and 16 million SNPs (Zhao et al. 2021). Han et al. (2022) also reported another FOV7-resistance QTL at a wide region (0.99-3.07 Mb) and its putative candidate gene on the same chromosome [*Gb_D03G0217*, named *GbCML* encoding for a calmodulin (CaM-like (CML) protein] identified in a *G. barbadense* RIL population of 110 lines through linkage mapping using 933,845 SNPs. Therefore, the terminal region of D03 carries a major resistance gene or QTL to both FOV4 and FOV7 in *G. barbadense*, but the exact location and the candidate genes remain to be resolved.

In Upland cotton, two major dominant resistance genes (*Fw_1_* and *Fw_2_*) to FOV7 were identified through a segregation analysis and an allelic test in China (Feng et al. 1998). A dominant resistance gene (designated *Fw^R^*, likely *Fw_1_* or *Fw_2_*) on D03 (c17) for FOV7 was later identified using 124 SSR markers in two F_2:3_ populations of Upland cotton (Wang et al. 2009). A major resistance QTL designated *Fov7* on D03 (likely *Fw^R^*) at 1.97-2.37 Mb was recently identified to be *Gh_D03G0209* (named *GhGLR4.8*) encoding a GLUTAMATE RECEPTOR-LIKE (GLR) protein through a GWAS of 290 Chinese Upland accessions using 2.7 million SNPs and confirmed using an F_2_ population and a CRISPR/Cas9-mediated knockout experiment (Liu et al. 2021). The resistance gene *GhGLR4.8* for *Fov7* in Upland cotton is apparently different from those candidate genes for FOV4 and FOV7 resistance identified in *G. barbadense*. However, it appears that both resistance genes/major QTLs for FOV4 and FOV7 in Upland cotton and *G. barbadense* are in a close proximity on the same chromosome D03, providing an incentive to target the chromosome region to develop markers for FOV4 resistance.

In the US, numerous Upland cotton accessions and germplasm lines have been evaluated for FOV4 resistance under the field and/or greenhouse conditions (Ulloa et al. 2006, 2020; Hutmacher et al. 2013; Abdelraheem et al. 2020; Zhang et al. 2020a, 2022d). Breeding lines and cultivars with improved FOV4 resistance have been developed and released through germplasm registrations in the public sector (Ulloa et al. 2009, 2016, 2022, 2023; Zhang et al. 2016, 2020a). However, no DNA markers associated with the resistance in these resistant lines and cultivars have been identified. Building on our previous studies in establishing a reliable screening method for FOV4 resistance in cotton (Zhang et al. 2020b, 2021, 2022a, b, c; Zhu et al. 2021b, 2022, 2023), the objective of this study was to develop the first set of SNP markers associated with FOV4 resistance in Upland cotton. The results will facilitate more molecular genetic studies which will accelerate the development of FOV4-resistant Upland cotton cultivars and cloning and isolation of the underlying resistance genes.

## Materials and Methods

### Materials

In this study, three sets of Upland cotton germplasms were used. Set 1 consisted of 223 Chinese Upland cotton lines and cultivars. This set of germplasm lines was used for evaluation of FOV4 resistance in the greenhouse (Fabian Garcia Plant Science Center, Las Cruces, NM) and development of resistance-associated markers. In this test, 15 seeds for each accession were sown in five hills (3 seed hill^-1^) in a 10-cm pot (as an experiment unit in each replication) filled with FOV4-infested potting soil from our previous studies (Zhang et al. 2020a). The germplasm lines were evaluated in 3 replications based on the method described in the following section.

Set 2 included 32 elite breeding lines from various US public cotton breeding programs and was used for marker analysis only based on results from Set 1. The entries were grown in a non-FOV4 infested field (Leyendecker Plant Science Center, New Mexico State University, Las Cruces, NM) in 2-rows × 7.62 m plots using a randomized completed block design with four replications. Seeds in Set 2 were mechanically planted in the field on May 2, 2022, and crop managements followed local recommendations.

Set 3 included 463 obsolete Upland accessions deposited in the US National Plant Germplasm System (NPGS). Seeds in this set were grown in the same greenhouse as Set 1. Seeds were sown in the same manner as Set 1 but in non-FOV4 infested potting soil. The commercial potting soil was Miracle-Gro Moisture Control Potting Mix 2 CF (Scotts Co., Marysville, OH, USA), which was either autoclaved or did not show any seedling diseases in negative control plants (i.e., without FOV4 inoculations) in our previous studies. To further confirm that it was free of cotton pathogens, the same source of commercial potting soil was used to grow six commercial cultivars (Acala 1517-08, Acala 1517-18 GLS, FM 2334GLT, PHY 725 RF, Pima DP 359 RF, and Pima PHY 881 RF). Plants without artificial inoculation of FOV4 did not show any FOV4-associated disease symptoms.

### Evaluation of FOV4 resistance

In Set 1, the FOV4 inoculum density in the infested soil was 6.8 × 10^5^ spores g^-1^ of soil. Seeds after sowing were covered with the same pathogen-free potting soil containing fertilizer as described above, followed by daily irrigation to water holding capacity. Wilted seedlings immediately after emergence subsequently died and were then scored, while a symptomatic or healthy plant after the 2-3 true leaf stage normally survived. Data on FOV4 related symptoms (seedling wilt and death) were recorded on a weekly basis until 21 days after planting (DAP). The disease incidence (DI, percentage of infected plants with symptoms) and seedling mortality (MR, percentage of dead plants) were calculated on a replication basis for each genotype.

To further confirm the resistance of the surviving seedlings, they were transplanted to a large pot (30 cm × 50 cm × 30 cm) containing FOV4-infested soil to grow to maturity. After boll harvesting, each plant was rated for root and stem vascular discoloration (RVD and SVD) based on a rating scale of 0 to 5 (Zhang et al. 2022c). Repotted plants that died were rated 5, while all other plants including a few chlorotic or necrotic or dwarf plants were rated for RVD and SVD. Seeds harvested from each plant were further used for a follow-up progeny test for FOV4 resistance to further reduce the chance of disease escapes, based on the method described above.

### Targeted SNP typing by sequencing using AgriPlex Genomics

Because of segregation in responses to FOV4 infections in most of the lines evaluated, leaf tissues were collected on an individual plant basis in Set 1 for DNA extraction using a quick cTAB method (Zhang and Stewart 2000). For Set 2, an unfolded leaf from each of five plants was collected and bulked to represent each line or cultivar. For Set 3, only one unfolded young leaf from each accession was harvested. DNA was quantified using a nanodrop spectrophotometer and adjusted to 20 ng μl^-1^.

Based on published QTLs for resistance to FOV (Said et al. 2015; www.cottonqtldb.org; Ulloa et al. 2013; Wang et al. 2018; Abdelraheem et al. 2017, 2020, 2021, 2022; Zhu et al. 2022), representative markers within consistent FOV-resistance QTL regions across two or more tests or publications were first examined in this study. As a result, a total of 75 flanking sequences for 75 SNPs targeting chromosome D03 spanning the 0.273-49.876 Mb region based on the sequenced TM-1 genome (HAU v1.1, Wang et al. 2019; www.cottongen.org) were chosen (Supplementary Table 1). DNA from the three sets of germplasm lines and cultivars were sent to AgriPlex Genomics (Cleveland, OH) for SNP typing based on multiplex PCR and sequencing. In brief, primers for these markers were synthesized in Integrated DNA Technologies Inc. (Coralville, Iowa). Primary multiplex PCR amplifications of the above targeted region were performed, followed by secondary PCR to add UDI indexes and Illumina tails. All amplicons were then pooled, purified, quantified, and diluted for sequencing using Illumina MiSeq. Sequences were analyzed for genotype calls and alleles reads using the PlexCall Data Analysis software (AgriPlex Genomics, Cleveland, OH).

### Statistical analysis

MR, SVD and RVD were calculated on a replication basis in Set 1. The analysis of variance was performed for the phenotypic results in FOV4 responses to calculate protected least significant difference (LSD) for a mean separation. Coefficients of correlation among the three evaluation parameters were also calculated on a genotype basis. Because of the limited number of SNPs on the targeted chromosome region of D03, a simple correlation analysis between SNPs and the phenotypic results was performed, followed by a t test to detect a significant difference between the two homozygous allele genotypes for the same SNP locus.

## Results and discussion

### Success rate in SNP typing using the AgriPlex Genomics platform

In this study, chromosome D03 was targeted using a total of 75 SNP markers. Among the 75 SNP markers designed, sequencing of the flanking regions for 46 SNPs using the AgriPlex Genomics platform was successful with a success rate of 61.33%. For the 46 successful SNPs, the mean sequence reads for each SNP in each DNA sample ranged between 108 and 5137 with an overall mean of 1,113 reads per SNP across all the SNPs for each DNA sample (Table 1). Therefore, the SNP calling for a DNA sample in this study was based on an average of 1,113 sequences amplified by a specific primer pair, and it was highly reliable. For the DNA samples representing the three sets of cotton lines, only 11.16% of the samples did not produce successful sequences (Table 1), indicating poor quality of DNA for these samples using the quick cTAB method without a cleanup procedure (Zhang and Stewart 2000). To ensure high quality with sufficient amount of DNA extracted from all cotton leaf samples, a cleanup procedure also based on cTAB (Zhang and Stewart 2000) should be implemented after the initial DNA extraction.

**Table 1.** Successful SNPs and alleles, mean reads, and percentage of successful DNA samples for each SNP using the AgriPlex Genomics platform.

| SNP Marker ID | Allele 1 | Allele 2 | No. mean reads | Successful samples (%) |
| --- | --- | --- | --- | --- |
| NMSU-FW001 | C | G | 690 | 86.49 |
| NMSU-FW003 | A | C | 550 | 83.17 |
| NMSU-FW006 | C | T | 2851 | 89.36 |
| NMSU-FW008 | C | T | 252 | 84.57 |
| NMSU-FW009 | C | T | 297 | 87.10 |
| NMSU-FW010 | C | T | 1585 | 91.80 |
| NMSU-FW012 | A | G | 1594 | 93.11 |
| NMSU-FW013 | A | G | 2163 | 93.90 |
| NMSU-FW015 | A | G | 979 | 89.54 |
| NMSU-FW016 | C | T | 1072 | 89.97 |
| NMSU-FW017 | G | T | 1594 | 93.55 |
| NMSU-FW018 | A | G | 704 | 92.24 |
| NMSU-FW019 | C | T | 1227 | 92.68 |
| NMSU-FW020 | A | G | 374 | 89.89 |
| NMSU-FW027 | A | G | 948 | 90.06 |
| NMSU-FW034 | C | T | 498 | 86.66 |
| NMSU-FW035 | A | G | 108 | 78.55 |
| NMSU-FW036 | A | C | 2242 | 89.71 |
| NMSU-FW037 | A | T | 629 | 79.08 |
| NMSU-FW039 | A | G | 432 | 91.02 |
| NMSU-FW041 | C | G | 2631 | 95.38 |
| NMSU-FW042 | A | C | 325 | 87.71 |
| NMSU-FW043 | A | G | 4137 | 91.89 |
| NMSU-FW045 | C | G | 1443 | 92.59 |
| NMSU-FW046 | C | G | 1387 | 92.07 |
| NMSU-FW047 | G | A | 476 | 90.15 |
| NMSU-FW048 | C | T | 1205 | 91.89 |
| NMSU-FW049 | C | T | 975 | 92.33 |
| NMSU-FW050 | A | T | 795 | 87.10 |
| NMSU-FW051 | A | G | 136 | 85.35 |
| NMSU-FW055 | A | T | 891 | 89.71 |
| NMSU-FW056 | A | T | 574 | 88.84 |
| NMSU-FW059 | A | C | 423 | 88.58 |
| NMSU-FW060 | A | G | 835 | 91.72 |
| NMSU-FW061 | C | T | 1822 | 92.76 |
| NMSU-FW069 | G | T | 2634 | 89.80 |
| NMSU-FW071 | G | A | 1917 | 93.46 |
| NMSU-FW072 | G | A | 175 | 83.35 |
| NMSU-FW073 | G | A | 1158 | 87.88 |
| NMSU-FW074 | A | T | 1471 | 90.06 |
| NMSU-FW077 | A | C | 1142 | 66.61 |
| NMSU-FW078 | T | G | 459 | 87.01 |
| NMSU-FW079 | G | A | 1603 | 91.89 |
| NMSU-FW080 | C | T | 659 | 91.54 |
| NMSU-FW082 | G | A | 234 | 83.61 |
| NMSU-FW083 | G | A | 909 | 90.93 |
| **Overall means** |  |  | **1113** | **88.84** |

Interestingly, each of the three pairs of SNPs (NMSU-FW021/NMSU-FW033, NMSU-FW005/NMSU-FW008, and NMSU-FW054/NMSU-FW055) was unintentionally designed from an identical sequence (Supplementary Table 1); however, only primers designed for NMSU-FW008 and NMSU-FW055 yielded positive PCR and sequence results. The same was true for each of the following two pairs of SNPs (NMSU-FW014/NMSU-FW015 and NMSU-FW043/NMSU-FW044), designed from the two different strands of the same sequence but with different lengths. Furthermore, two SNPs were successfully sequenced for three SNPs (NMSU-FW036/NMSU-FW040/NMSU-FW069) that were also unintentionally designed from the two different strands of different lengths from the same DNA sequence (Supplementary Table 1). The multiplex PCR and sequencing produced the same sequence with the same SNP calling for the complementary strands (A/C vs. T/G). The results indicate high accuracy of sequencing and high specificity of the two different primer pairs targeting the same sequence and SNP. Therefore, for must-have key SNPs, it is always a good practice to design more than one pair of primers targeting the same SNP but amplifying different lengths of the flanking sequences.

### SNP markers associated with FOV4 resistance

Among the three evaluation parameters in Set 1, MR was significantly but weakly correlated with SVD (r= 0.291, *P*<0.01) and RVD (r= 0.169, *P*<0.05); however, SVD and RVD was highly significantly correlated (r= 0.712, *P*<0.0000), consistent with our previous reports (Zhang et al. 2022b; Zhu et al. 2023b). In this test, ANOVA detected significant differences in responses (MR, SVD and RVD) to FOV4 infections, warranting a further analysis between the three evaluation parameters and SNP markers. The progeny test for the seeds harvested from survived plants grown in FOV4-infested soil confirmed the results from the three replications of the test. Once adequate seeds from these germplasm lines are obtained, they will be further evaluated for FOV4 resistance under field (Fabens, Texas) conditions and made publicly available. Based on a set of eight cotton cultivars, we (Zhang et al. 2022b, c, f) previously demonstrated that evaluation results for FOV4 resistance in the greenhouse were consistent with these under the field (Fabens, Texas) conditions, but with a lower experimental errors.

For the 46 successful SNP markers, a simple correlation analysis was performed between the SNPs and MR in Set 1. No significant correlation was detected for any SNP (Fig. 1), suggesting that the chromosome region as anchored by the SNPs is not associated with MR. However, 24 SNPs in the region (2.13-2.29 Mb) were significantly correlated with vascular discoloration which peaked at 2.249 Mb, and the correlations of the SNPs with SVD were similar to these of the same SNPs with RVD (Fig. 1). The result indicates that the region on D03 carries gene(s) conferring resistance against both RVD and SVD caused by FOV4.

**Fig. 1.**
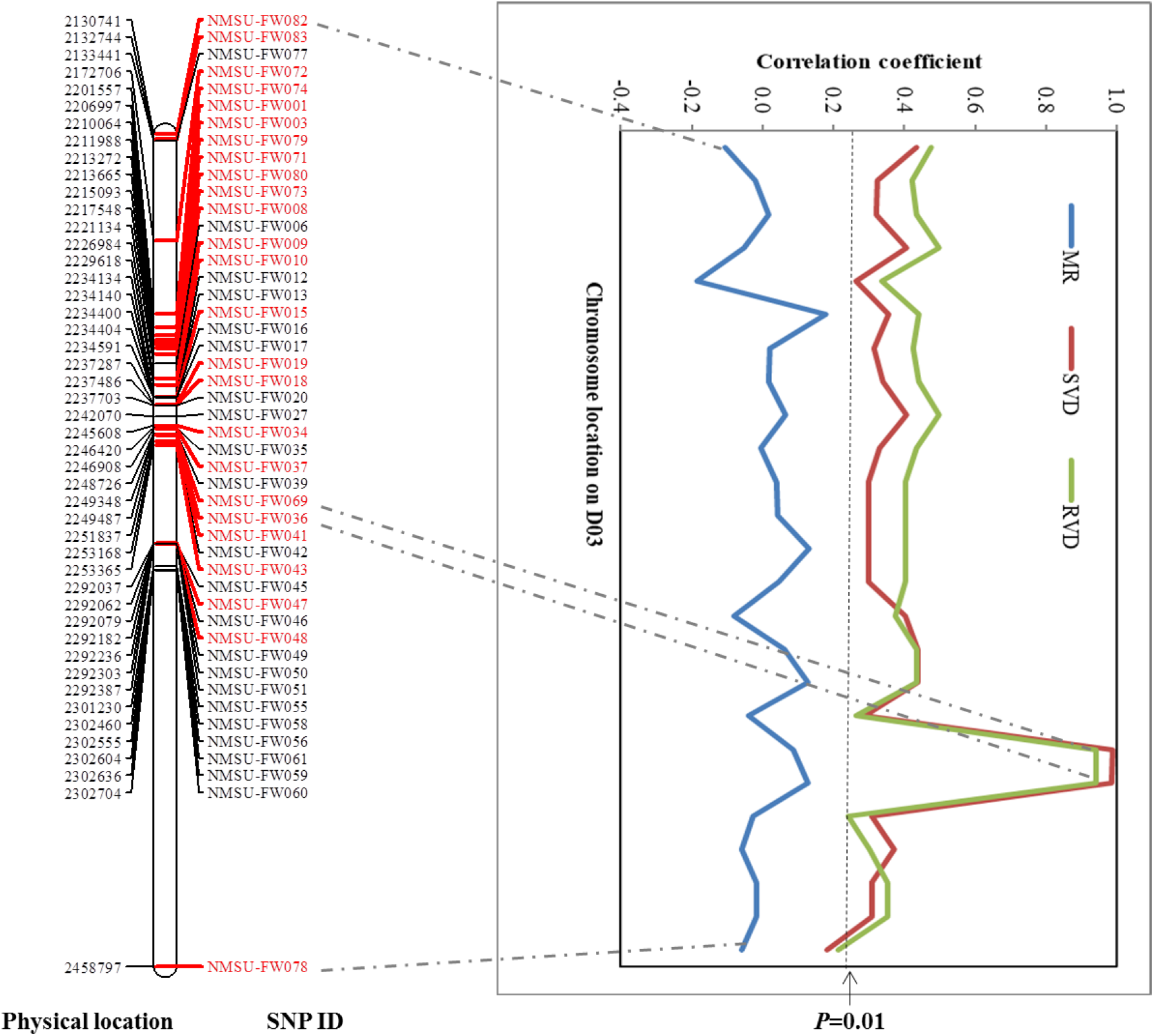
Chromosome location of SNPs on D03 and their correlation with mortality rate (MR), stem vascular discoloration (SVD) and root vascular discoloration (RVD) in 223 Chinese Upland cotton accessions. SNP ID in red color showed significant correlation with SVD and RVD.

A further analysis was performed using the two most significant (peak) SNP markers (NMSU-FW036 and NMSU-FW069) in Set 1. The genotypes with homozygous A or T SNP allele averaged significantly lower mean SVD (0.88 vs. 2.54) and RVD (1.46 vs. 3.02) than the genotypes with the homozygous C or G SNP allele (Fig. 2). Although the AA SNP genotypes averaged a lower mean MR (41.82 vs. 50.88%) than the CC SNP genotypes, the different was not significant. This was evidenced from our observations that numerous dead plants possessed the A SNP allele, while other survived plants but with SVD and RVD had the C SNP allele.

**Fig. 2.**
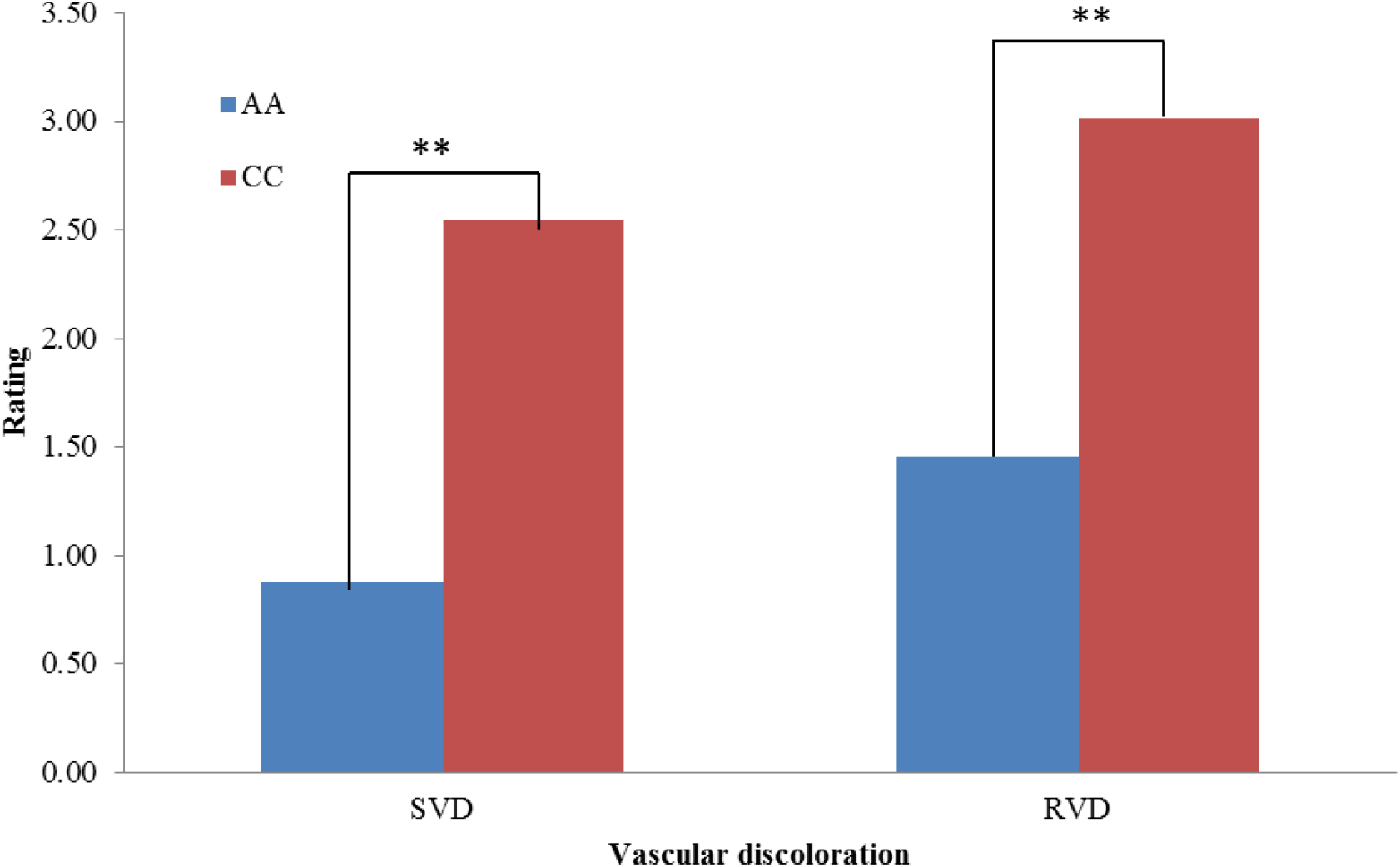
Differences in mean stem vascular discoloration (SVD) and root vascular discoloration (RVD) between genotypes with A or T and C or G alleles in 223 Chinese Upland cotton accessions. \*\**P*<0.01.

### FOV4 resistance-associated marker analysis within US and Chinese Upland cotton

The two most significant SNP markers (NMSU-FW036 and NMSU-FW069) were used to compare frequencies among different groups of Upland cotton germplasm lines (Table 2). The results showed that 37.22% of the Chinese Upland cotton cultivars and lines tested (Set 1) were homozygous in possessing the diagnostic SNP markers for FOV4 resistance and 11.66% of them were heterozygous in the SNPs. However, none of the 32 elite public breeding lines (Set 2) carried the FOV4 resistance-associated A allele SNP markers. Interestingly, four (0.86%) of the 463 obsolete US Upland accessions (Set 3) assayed possessed the SNP A allele for FOV4 resistance including two in a heterozygous status. Therefore, resistance to FOV4 does exist within the US Upland cotton germplasms. Because all the US accessions were developed under non-FOV4 field conditions, it is understandable that FOV4-resistant genotypes were randomly fixed in breeding lines and cultivars at a very low frequency. However, selection for FOV4 resistance will undoubtedly increase the frequency of resistance genes and genotypes. For example, China has had numerous extensive breeding programs for resistance to FOV7 since the 1970s, and it in fact requires such a resistance before a cultivar is approved for production at the provincial or central government level (Zhang et al. 2015; Zhang 2018). Liu et al. (2021) showed that the frequency of the FOV7-resistance gene (*GhGLR4.8^A^*) allele genotype was as high as 43% in 289 Chinese Upland cotton cultivars, as compared to only 7% among 32 introduced Upland accessions from the US, India, and Brazil.

**Table 2.**
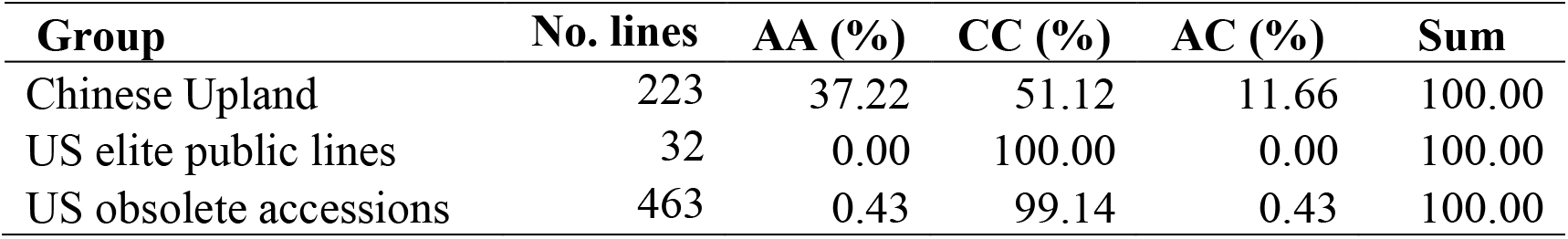
The frequency of a diagnostic SNP NMSU-FW036 (A/C) within different groups of Upland cotton lines.

However, whether the 48.77% FOV4-resistance allele-carrying Chinese Upland lines and cultivars identified in this study are also resistant to FOV7 is currently unknown and should be studied. Because the flanking sequences for the two SNPs (NMSU-FW036 and NMSU-FW069) are identical to the resistance gene (*Fov7*) *GhGLR4.8^A^*, it appears that the chromosome region carrying *GhGLR4.8^A^* with the A SNP allele confers resistance to vascular discoloration caused by both FOV4 and FOV7 through root infections. However, it does not provide resistance against seedling damping-off (mortality) from hypocotyl infection by FOV4. Unlike FOV4, FOV7 under field conditions normally does not cause seedling mortality (Zhang et al. 2022e). Because of the uniqueness of FOV4, resistance to seedling damping-off caused by FOV4 requires a different genetic mechanism in cotton. Previously, we showed that seedling mortality caused by FOV4 was correlated with SVD and RVD, but the associations were far from perfect (Zhang et al. 2022b; Zhu et al. 2023b), suggesting both common and independent resistant mechanisms for MR and vascular discoloration.

## Conclusions

Although numerous QTLs associated with FOV4/FOV7 resistance have been reported, no diagnostic markers for a major resistance gene or QTL was previously reported and used by breeders to develop FOV4/FOV7-resistant Upland cotton cultivars. For the first time, this study has developed a set of SNP markers (through genotyping by targeted sequencing) for selecting Upland cotton for resistance to FOV4 and has identified many FOV4-resistant Upland cotton germplasms with the diagnostic markers for the resistance. The SNP calling for each cotton line was based on an average of 1,113 sequence reads and was therefore highly reliable, and the consistency was further validated from a total of 719 cotton lines using the targeted sequencing technology in this study. The flanking sequences for the SNPs provided in this study will be useful for the cotton community to design primers in developing other SNP typing methods such as polyacrylamide gel-based single-strand conformational polymorphisms (SSCP), TaqMan, and KASP (Broccanello et al. 2018). The diagnostic SNP markers developed and their related sequences will be highly useful for others to genotype germplasm lines and advanced breeding materials for FOV4 resistance. For the first time, the results have further demonstrated that the same chromosome region carrying FOV7 resistance gene (*Fov7*-*GhGLR4.8^A^*, Liu et al. 2021) on D03 also confers resistance to FOV4 in Upland cotton. However, whether the resistance genes for FOV4 and FOV7 are identical, allelic or closely linked is currently unknown. There may be other genes residing in other genome regions, requiring a genome-wide association study (GWAS) using genome-wide high density markers or linkage mapping using segregating populations from parents with different sources of resistance.

